# Traumatic Brain Injury induces persistent behavioural deficits that are rescued by the antiepileptic drug Levetiracetam

**DOI:** 10.64898/2026.09.12.751143

**Authors:** Shayleen Ghassemi, Xavier Penetrante, Tanja Zerulla, Trevor J. Hamilton, W. Ted Allison

## Abstract

Traumatic Brain Injury (TBI) produces acute challenges including seizures and neuroinflammation, and these then exacerbate lifelong difficulties including epilepsy, dementias, and anxiety. Applying antiepileptic drugs (AED) to manage acute seizures after TBI has promise to slow these progressive deficits. Levetiracetam (LEV) is an AED that mitigates seizures and neuroinflammation in various contexts. Here, we explore the utility of the larval zebrafish TBI model by documenting long term behavioural deficits (if any) induced by larval neurotrauma and whether they can be mitigated by LEV. We induced a mild blast TBI in three days-post-fertilization larvae, administered LEV (3 × 10^-2^ mM) for two days, and assessed behaviour in adults (9±1 or 22±1 months later). In open field tests, adult zebrafish that experienced TBI as larvae exhibited increased anxiety-like behaviour compared to sham-treated larvae, but this resolved by later adulthood. LEV treatment immediately following larval TBI did not significantly alter these outcomes. Novel object approach tests revealed sustained deficits in exploratory behaviour: adults that experienced larval TBI spent significantly less time investigating a novel object compared to sham controls throughout adulthood; this suggests long-term alterations in fear-related cognitive processing or threat evaluation. LEV treatment increased time spent near the novel object in early adulthood, but not later adulthood, suggesting a time-dependent efficacy of early pharmacological intervention. In sum, the larval zebrafish TBI model is suitable for investigating the progression and treatment of behavioural deficits. LEV showed only limited efficacy, but adjusted AED regimens have potential to mitigate long-term behavioural outcomes of TBI.

**Highlights:**

- Blast TBI on larval zebrafish provides a bioethically-favorable neurotrauma model
- TBI on larval zebrafish produces subtle but enduring behavioural deficits
- The antiepileptic Levetiracetam had a modest ability to mitigate deficits
- Applied briefly after TBI, Levetiracetam mitigated anxiety-like deficits in adults

## 1. Introduction

TBI induces a complex suite of neuropsychological problems and a heightened risk of dementia (Mavroudis et al., 2024; Scheenen et al., 2017). The progressive nature of TBI outcomes is increasingly emphasized as a priority issue in clinical and experimental models (Katz et al., 2021; Maas et al., 2022). The quality and progression of deficits can be different after pediatric vs. adult TBI, and a better appreciation of chronic outcomes following TBI is needed (Kurowski, Haarbauer-Krupa, & Giza, 2023). Understanding the complexity of disease progression after neurotrauma, and evaluating treatments, is hampered by a limited range of model systems. Cell cultures, organoids and invertebrates are of limited utility for investigating TBI neurotrauma, and rodent TBI models are resource-intensive and face ethical barriers that may hamper exploration and translation of promising ideas or results.

A recently introduced preclinical model of TBI uses larval zebrafish to fill the gap in available neurotrauma platforms and provides a bioethically-favourable model for TBI research (Alyenbaawi et al., 2021). It applies a repeated mild blast injury where larval fish in water are secured within a sealed syringe and uses controlled hits to the syringe plunger to drive pressure waves that damage the CNS and other organs. The injury is most akin to TBI experienced by victims who were in the vicinity of an explosion (Nakagawa et al., 2011). The advantages of a larval zebrafish model include the lower risk/investment associated with exploring how compounds and polypharmacy strategies alter neurological outcomes. Zebrafish larvae provide high-throughput, genetic tractability and transparent tissues enabling observation of antemortem dementia biomarkers. TBI in larval zebrafish induces events in the subsequent days that are akin to events in TBI patients and rodent models: post-traumatic-seizures, reduced blood flow, CNS cell death, tau aggregation, and neuroinflammation (Alkhatib, Zerulla, Finlay, & Allison, 2026; Alyenbaawi et al., 2021; Locskai, Ghassemi, Tan, Kinley, & Allison, 2026). These various endpoints are all suitable for high-throughput *in vivo* testing of TBI pathomechanisms and treatments. However, the larval zebrafish TBI model lacks metrics of cognition that are essential outcome measures in animal models of neurotrauma and dementia. Broadly speaking, larval zebrafish are not reliable for measuring cognitive deficits in high-throughput behavioural paradigms, and so here we sought to determine if behavioural deficits were observable in adult zebrafish that had received TBI as larvae.

Beyond its impacts on cognition, TBI is also associated with profound and persistent neurobehavioural alterations in anxiety and fear responses. In humans, anxiety disorders following TBI are prevalent, with reported rates ranging from 11–70%, substantially exceeding the prevalence in the general population (Jorge & Arciniegas, 2014). Even ‘mild’ TBI can increase the prevalence of post-traumatic stress disorder (PTSD) to 19% at 6 months post-injury (Glenn et al., 2017; Stein et al., 2019). TBI in rodents leads to increased avoidance and anxiety-like behaviour in open field and elevated maze tests, with effects that can vary by test and post-injury time point (Tucker & McCabe, 2021). Mild TBI also induces changes in limbic circuitry that manifest as increased conditioned fear and emotional dysregulation, consistent with PTSD-like phenotypes (Meyer, Davies, Barr, Manzerra, & Forster, 2012). Pertaining to our analysis, anxiety and fear represent related but distinct behavioural domains: anxiety reflects sustained responses to diffuse or uncertain threats, whereas fear responses are elicited by discrete stimuli. Although zebrafish have been widely used to model PTSD-like symptoms through measurable behaviours such as thigmotaxis and reduced exploration (Al-Zoubi et al., 2024), these domains remain underexplored in zebrafish models of TBI.

The timeline of behavioural deficits following TBI (if any) may be predicted to be progressive, considering the inexorable and progressive nature of dementias (e.g. AD and CTE) for which TBI is a cause or leading risk factor (Mavroudis et al., 2024). However, predicting a timeline of behavioural deficits may also consider that zebrafish have substantial capacity for CNS regeneration during development and in adulthood (Fleisch, Fraser, & Allison, 2011; Pose-Mendez et al., 2023; Zambusi & Ninkovic, 2020). In fish, brain repair might be imagined to mask or eliminate trauma-induced long-term behavioural deficits. The outcomes of the current study were difficult to predict.

Thus, our goal was to expand the utility of larval zebrafish as a preclinical model for neurotraumas, especially the capacity to address cognition and anxiety. In parallel, we sought to test levetiracetam (LEV, an anti-epileptic), applied briefly after TBI, as a potential prophylactic to mitigate any behavioural and/or cognitive deficits. LEV acts in the presynaptic compartment by binding synaptic vesicle protein SV2a to modulate neurotransmitter release which reduces aberrant high frequency activity (Lynch et al., 2004; Y. Zhang et al., 2022). LEV has shown promise in clinical trials for slowing dementia progression (Sen et al., 2024; Vossel et al., 2021), consistent with a growing literature noting bi-directional causal links between epilepsy and neurodegenerative disease (Liu & Barr, 2023; Locskai, Alyenbaawi, & Allison, 2024; Mohamadi et al., 2025; Sen et al., 2026; D. Stewart & Johnson, 2025). LEV may also have non-canonical actions on neuroinflammation, e.g. via modulating *Fosl1* expression and suppressing microglia (Komori et al., 2022; Niidome et al., 2021; Y. Y. Zhang et al., 2023).

We recently showed that LEV, applied briefly following TBI in larval zebrafish, was able to prevent CNS cell death and tau aggregation when measured days later (Locskai et al., 2026). Indeed, the larval TBI model shows AEDs reduce, and convulsants increase, TBI-induced tauopathy, cell death and neuroinflammation (Alkhatib et al., 2026; Alyenbaawi et al., 2021; Locskai et al., 2026). These trials have focussed on LEV and retigabine (RTG), applied for two days following TBI (the protocol we mimicked here) and this reduces deficits, often to near-baseline levels, when assessed two days after that. In rodents, brief application of RTG after TBI was able to reduce behavioural deficits months later (Vigil et al., 2023; Vigil, Carver, & Shapiro, 2020). Here, we selected LEV due to its ongoing clinical development towards slowing cognitive decline, and for its demonstrated promise as a component of polypharmacy approaches (Caudle, Lu, Mountney, Shear, & Tortella, 2016; Locskai et al., 2026; Metcalf et al., 2017; Zou et al., 2013).

Our original goal was to evaluate the long-term impacts of TBI (and LEV treatment) on behavioural measures of cognition, while also measuring anxiety-like behaviour metrics as an essential element in interpreting the outcomes. The results suggest that TBI on larval zebrafish produces long-lasting impacts on anxiety-like behaviour levels. In contrast, our assessments of cognition using novel object recognition tests were equivocal, but this represented a technical (rather than biological) challenge because even our control sham animals failed to demonstrate the expected levels/consistency in learning and memory tasks; we note that stress is often a complicating confound for interpreting outcomes of these novel object recognition tasks (Gaspary, Reolon, Gusso, & Bonan, 2018; Hamilton, 2018). Thus, we focus our interpretation on the chronic consequences of TBI as it pertains to anxiety-like behaviours. The data suggest that LEV, applied briefly after TBI in larval zebrafish, can mitigate these anxiety-like deficits, at least at the single dose and particular time-course of LEV we trialed here.

## 2. Methodology

### 2.1 Zebrafish Husbandry

Wildtype fish (AB strain) were raised and maintained within the University of Alberta fish facility under a 14/10 light/dark cycle from 8:00 am to 10:00 pm at 28±1°C as previously described (Westerfield, 2000). All procedures for care of adult zebrafish were approved by the Animal Care and Use Committee: BioSciences at the University of Alberta, operating within the guidelines of the Canadian Council on Animal Care. This research adhered to ARRIVE guidelines for animal research.

### 2.2 Traumatic Brain Injury

In this study, Alyenbaawi’s model of TBI was used to induce TBI (Alyenbaawi et al., 2021; Gill et al., 2024; Locskai et al., 2025). Approximately 10-15 larval zebrafish at 3 days-post-fertilization were placed into a 20mL syringe (Becton Dickinson #302830) with 1 mL of E3 media. The syringe was closed with a Lok stopper valve attachment and placed in a three-pronged clamp, with the syringe plunger centered under a 1.08 m guide tube (Figure 1A). A 200 g weight was dropped into the guide tube, creating a shock wave through the liquid media as it hit the plunger. The shock wave caused by the impact of the weight induced brain trauma. The details of the TBI methods and configuration (size of syringe, height of tube, and weight) were selected based on previous findings that showed the optimal combinations for behaviour disparities on larvae while retaining minimal impacts on survival or morphology (Locskai et al., 2025). This configuration delivers a maximal pressure wave of about 300 kPa (Locskai et al., 2025). The weight was dropped 3 times on each sample, and the larvae were repositioned between each drop, to mitigate inter-individual variability in brain injury. The sham procedure involved placing the larvae in the syringe and following all the previously noted steps except the weight drop.

**Figure 1.**
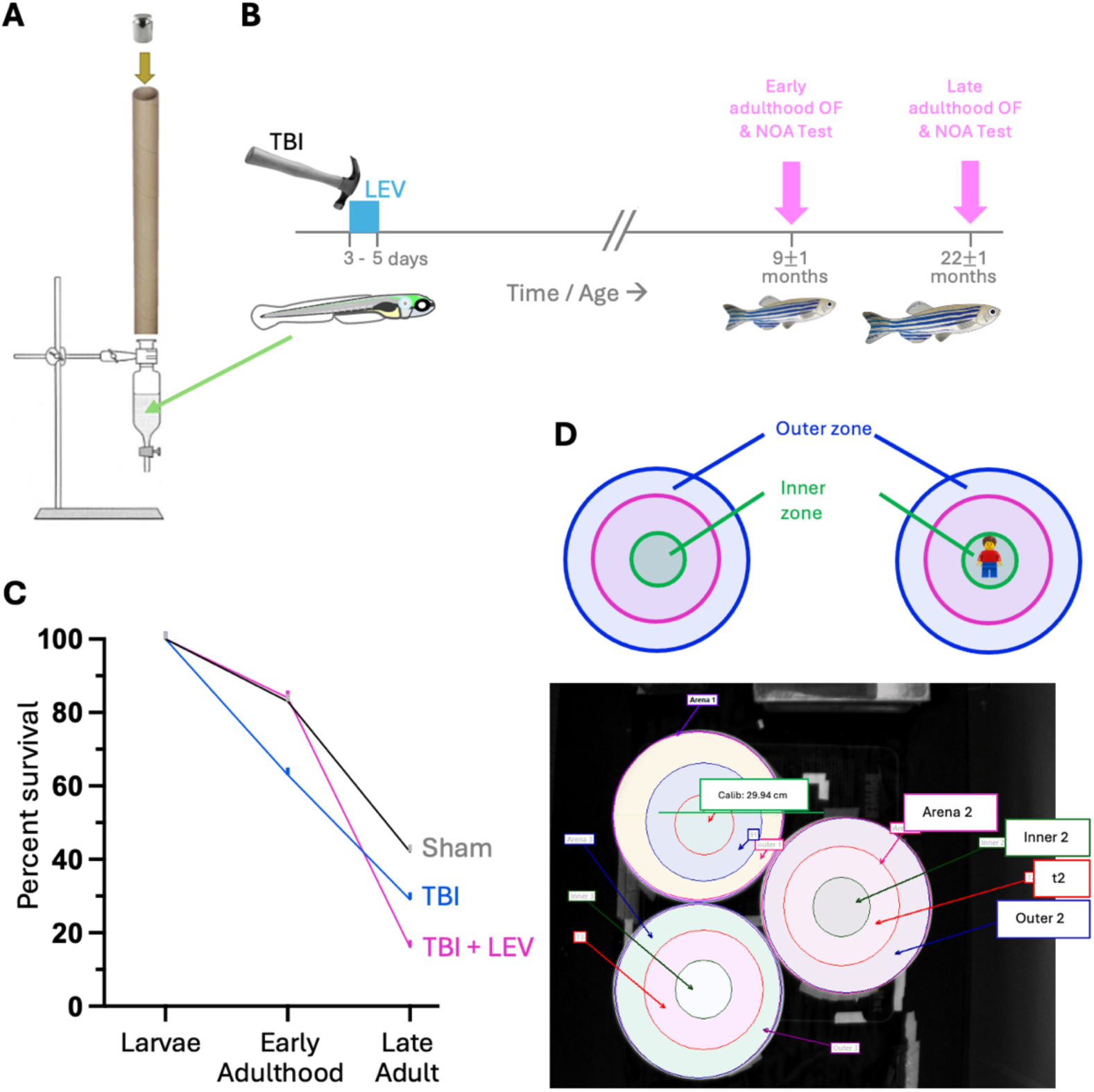
Treatment administration (injury and drug) and behavioural testing. **A)** Schematic representation of the apparatus used to induce traumatic brain injury (TBI) in larval zebrafish. As described by Alyenbaawi et al. (2021), the setup consists of a 1.08 m-tall tube with a syringe plunger inserted at one end, accommodating approximately 10–15 zebrafish in 1 mL of fish media. A 200 g weight was released onto the plunger of the sealed syringe, generating a shock wave that induced brain trauma. **B)** The experimental timeline begins with induction of traumatic brain injury (TBI) followed immediately by two days of treatment with the anti-epileptic drug (AED) Levetiracetam (LEV at 10^-2^ mM), with two behavioural experiments conducted at two ages in adulthood: open-field (OF) and novel object approach (NOA). **C)** Survival trends between experimental treatment groups, displayed as number of zebrafish in cohort 1 across each time point. Sample number at larval stage; Sham n= 36, TBI n= 38, TBI + LEV n= 38. Sample number at early adulthood; Sham n= 30, TBI n= 24, TBI + LEV n= 32. Sample number in late adulthood; Sham n= 15, TBI n= 11, TBI + LEV n= 6; No differences in survival were identified (*F*(2, 4) = 0.93, *P* = .4633). **D)** Set up for open field-testing arena without an object and the novel object approach test with a multicolored Lego figurine affixed to the center of the open field arena. There are three virtual zones divided equally based on the diameter (inner, transition, and thigmotaxis zones).

### 2.3 Drug administration and removal

Immediately following TBI, approximately half of the sham and TBI groups were exposed to either E3 medium or Levetiracetam (3 × 10^-2^ mM LEV, Millipore Sigma, product number L8668) dissolved in E3. The drug dosage was selected based on a previous study that investigated the efficacy of LEV doses post-TBI (Locskai et al., 2026). The drug was washed out via a media change at 48 hours-post-injury. The new media consisted only of E3. The four groups consisted of (i) Sham + no LEV treatment, (ii) Sham + LEV treatment (only in cohort 2), (iii) TBI + no LEV treatment, and (iv) TBI + LEV treatment.

### 2.4 Behavioural Testing

Following TBI and drug administration, fish were placed back into the animal habitat and reared under identical conditions. Adult zebrafish were tested in an open field test (OFT) followed by a novel object approach test (NOA), respectively, beginning when the zebrafish reached 9 months of age (Figure 1B). Fish were tested longitudinally at two experimental time points, early adulthood (9 ± 1 months) and late adulthood (22 ± 1 months). Both adulthood stages included two independent cohorts. *Cohort one* included three experimental groups; sham, TBI, TBI + LEV, and the sex of the animals was documented only at the late adulthood stage. *Cohort two* assessed three experimental groups at early adulthood (sham, TBI, TBI + LEV), and four experimental groups at late adulthood (sham, TBI, TBI + LEV, sham + LEV), and sex was documented at both adulthood stages. The addition of the sham + LEV group was to determine whether LEV is inducing any defects on its own. The adult behavioural tests were conducted from 11:00 AM to 6:00 PM in accordance with the middle hours of the light cycle (8:00am – 10:00pm). Sample sizes for cohort 1: Early adulthood= Sham n= 30, TBI n= 24, TBI + LEV n= 32. Sample sizes in late adulthood; Sham n= 15, TBI n= 11, TBI + LEV n= 6. cohort 2: early adulthood: sham: 20, TBI= 14, TBI + lev= 16, Late adulthood: sham = 18, sham + lev= 25, TBI= 11, TBI + LEV= 12.

#### 2.4.1 Open field test on adult zebrafish

To evaluate zebrafish anxiety-like behaviour and movement, the open-field test paradigm was employed, as described in previous studies (Johnson, Verbitsky, Hudson, Dean, & Hamilton, 2023; Scatterty & Hamilton, 2024; A. Stewart et al., 2012). Individual adult zebrafish were each placed individually in a circular arena (diameter: 26 cm) filled with 5 of habitat water (refreshed for each fish). The time spent in three concentric virtual zones (thigmotaxis, transition, and inner zones) were quantified in Noldus EthoVision XT, v.17). The inner zone had a radius of 4.3 cm, the transition zone spanned from the inner zone to the next concentric circle with a radius of 8.6 cm, and the thigmotaxis zone covered the area from the transition zone to the wall of the arena (Figure 1D). Anxiety-like behaviour was inferred by fish spending more time in the thigmotaxis zone, close to the walls of the arena (Figure 1D). More time spent in the inner zone also denoted a reduction in anxiety-like behaviour (A. Stewart et al., 2012). Testing occurred in white circular plastic arenas with a diameter of 26 cm, and a height of 16 cm. Temperature was maintained at 26 ± 1 °C. For each trial, the arenas were filled to a height of 6 cm with fresh habitat water and were replaced every four trials. Immobility and total distance moved were measured to quantify locomotor disruptions affecting exploratory behaviour in each group. The threshold for immobility was set to a 5% in EthoVision, such that a lack of movement is denoted when there is less than 5% change in the pixels between videoframes (section 12.3.3 of EthoVision 3.1 Reference Manual).

#### 2.4.2 Novel object approach test on adult zebrafish

The novel object approach (NOA) test is another validated behavioural assay employed to evaluate anxiety-like behaviour and fear response towards a specific object (A. Stewart et al., 2012). Fear responses are intimately tied to cognitive processes, encompassing perception, interpretation, memory, learning, and executive functions (Adolphs, 2013). Cognitive mechanisms enable individuals to perceive and interpret threatening stimuli, form fear memories, and exert control over fear responses. Following an OFT test trial, a LEGO® figurine (∼4.3 cm tall), constructed with multiple colors to mitigate colour bias, was positioned at the center of the arena (Figure 1D). Motion-tracking recording via EthoVision was initiated immediately. Each trial lasted for 10 minutes. The parameters measured in each trial included distance moved, immobility and time spent in thigmotaxis, transitions, and inner zones (as described above).

### 2.5 Statistical Analyses

Statistical analyses were performed using GraphPad Prism version 10. Data were assessed for normality using the D′Agostino-Pearson omnibus normality test. Data were analyzed using an ordinary one-way ANOVA followed by post-hoc Dunnett’s multiple comparison test. Non-parametric data was analyzed using a Kruskal-Wallis with post-hoc Dunn’s multiple comparison test. Any fish that did not move for the entirety of the experiment were omitted from statistical analysis.

## 3. Results

### 3.1 Survival Trends Across Groups receiving TBI and/or LEV

The fish were treated with TBI or a sham procedure as larvae and characterized at multiple stages of adulthood. Fish were assessed when they were at 9±1 and 22±1 months old, which we denoted as “early” and “late” adulthood to simplify the discussion of outcomes. Note, however, that zebrafish reach adulthood (sexual maturity) by about 3 months of age, and often thrive to be 4 or 5 years old (though they are typically euthanized at 2 years old in accordance with standard operating procedures). Thus, our groups of fish were not very young adults, nor were they old adults, but allow a comparison of outcomes over a relatively large and informative timespan that aligns with practicalities of most zebrafish research colonies.

An immediate challenge of the study design, when exploring to identify adult behavioural deficits in a new larval TBI model, was selecting an appropriate level of injury. One predicts that a weak injury would produce false negative outcomes, where the deficits were too mild to detect amongst the inter-individual variation. At the opposite extreme, an injury of too great an intensity may produce comorbidities and mortalities that prevent interpretable outcomes. A relatively mild injury dose was selected, based upon the behavioural and pathological outcomes observed at various injury doses 4 days post-injury (Locskai et al., 2025). The number of fish at the larval stage, early adulthood, and late adulthood for cohort 1 was tracked and no significant difference in the survival trend was observed among treatments (Figure 1C) (*F* (2, 4) = 0.93, *P* = .4633). The fish from each treatment were not significantly different in their size (standard length) at the time of euthanasia.

### 3.2 Locomotion-Open Field and Novel Object Approach Test

Locomotor activity was assessed during both the open field (OFT) and novel object approach (NOA) tests to determine whether traumatic brain injury (TBI) or levetiracetam (LEV) treatment altered movement patterns. Across all experimental groups (Figure 2A-H), the only significant difference was observed in total distance moved in NOA test at late adulthood (*H* (4) =9.189, Figure 2H) with post-hoc differences between injured and non-injured fish that had been administered LEV (*P* = .0269). LEV-administered fish moved a longer distance when injured compared to their uninjured counterparts. These findings suggest that LEV treatment may affect immobility outcomes locomotor behaviour in the fish but only after brain injury.

**Figure 2.**
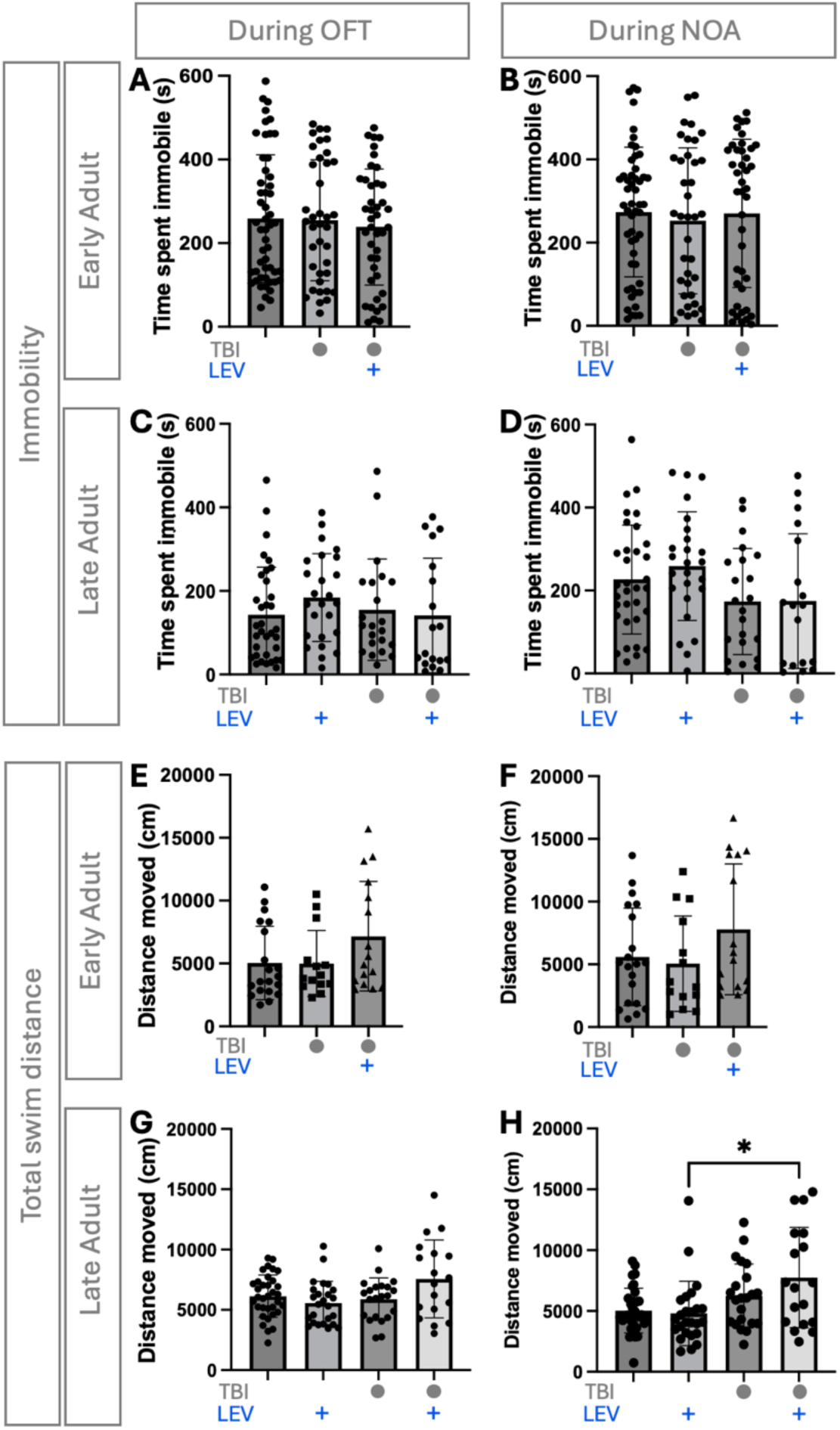
Locomotion during the behavioural tests. A) Time spent immobile during open field tests (OFT) in early adulthood (*P* = .8531). B) Time spent immobile in novel object approach (NOA) test at early adulthood (*P* = .85705). C) Time spent immobile in OFT at late adulthood (*P* = .2274). D) Time spent immobile in NOA test at late adulthood (*F*(3.000, 74.62) = 2.043, *P* = .1132). E) Total distance moved in OFT at early adulthood (*F*(2.000, 36.71) = 2.167, *P* = .1290). F) Total distance moved in NOA test at Early adulthood (*P* = .1583). G) Total Distance moved in OFT at late adulthood (*F*(3.000, 48.14) = 3.002, *P*= 0.1356). H) Total distance moved in NOA test at late adulthood (*P*= 0.0269). The bars indicate the mean of each group with SEM. Each data point represents an individual zebrafish. Levetiracetam (LEV) was applied at 10^-2^ mM as per the timings in Figure 1B, immediately after TBI in larval fish and months prior to these behaviour recordings.

### 3.3 Anxiety-like Behaviour in the Open Field Test (OFT)

The OFT evaluated zebrafish anxiety-like behaviour which was quantified by the amount of time spent in the thigmotaxis zone, close to the walls of the arena (Figure 1D). Adult fish that had undergone TBI at 3 days post-fertilization spent a significantly higher amount of time in the thigmotaxis zone (*H* (3) = 9.728, *P=*.0077) (Figure 3C) with post-hoc differences between the sham group and TBI group,(*P=*0.0203) as well as the sham group and TBI + Lev group (*P*= 0.0298). The results indicate an increase in anxiety in zebrafish following TBI that is not rescued by LEV treatment. At late adulthood, there was no significant difference in time spent in thigmotaxis zones between any groups (H (4) = 4.755*,P=* .1907, Figure 3D), this could be attributed to an age-related increase in baseline anxiety levels.

**Figure 3.**
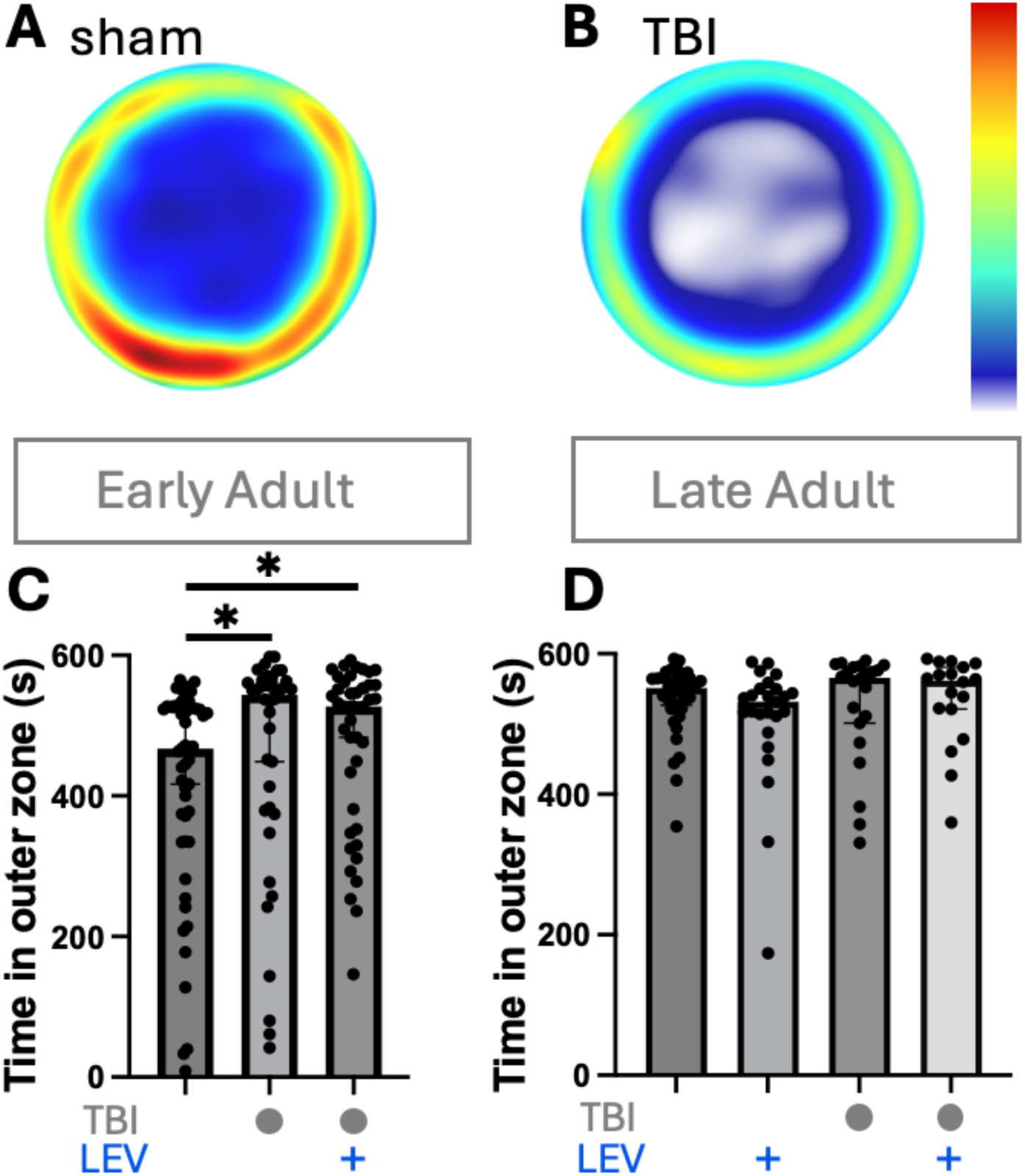
Anxiogenic effects of traumatic brain injury (TBI) in open field tests measured by time spent in the outer zone (thigmotaxis). Levetiracetam (LEV) was applied at 10^-2^ mM as per the timings in Figure 1B, immediately after TBI in larval fish and months prior to these behaviour recordings. **A, B)** Activity of an exemplar fish from the sham and TBI groups, showing the latter exhibited increased time spent in outer zone, represented as heatmaps of the circular arena. The colours denote amount of time spent in each location, with hotter/red colours indicating longer duration. **C)** Thigmotaxis in early adulthood (*P= .0077*) (Sham n= 50, TBI n= 37, TBI + LEV n= 46). **D)** Thigmotaxis in late adulthood (*P=* .01907) (Sham n= 32, Sham + LEV n= 21, TBI n= 22, TBI + LEV n= 18). Each data point represents one fish. The bar indicates the median of each group with 95% Cl. Significant differences between groups are indicated by (*): *P*< .05.

### 3.4 Behavioural Deficits in Novel Object Approach Test

At early adulthood, TBI-administered fish spent significantly less time in the inner zone exploring the novel object (*H* (3) = 24.93, *P* <.0001, Figure 4D),with post-hoc differences between the sham and TBI group (*P*<.0001) and TBI and TBI + LEV group (*P=* 0.0165). LEV treatment, therefore, decreased TBI-induced behavioural deficit at early adulthood (Figure 4D).

**Figure 4.**
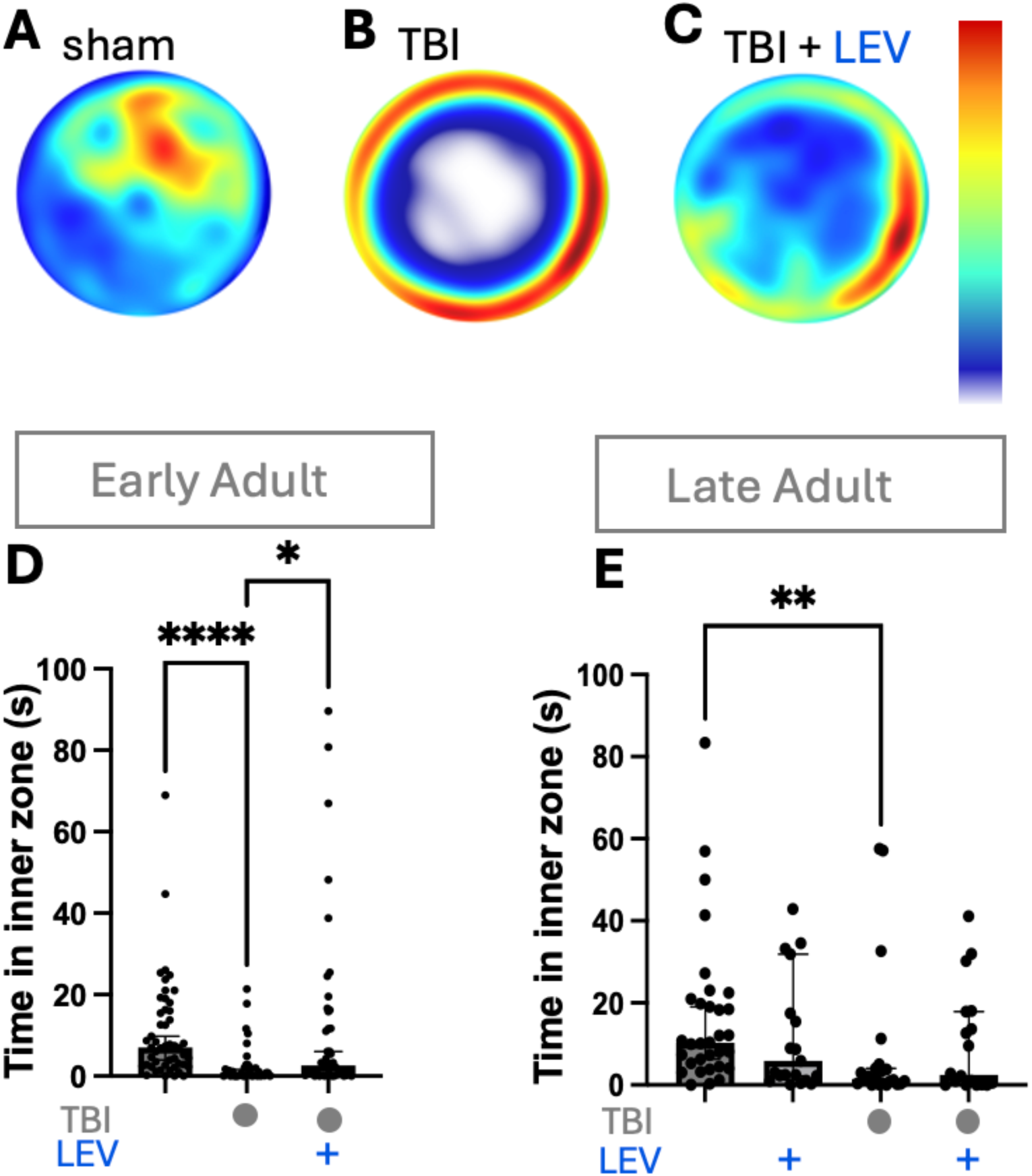
Boldness-related effects of traumatic brain injury (TBI) in novel object approach tests. Levetiracetam (LEV) was applied at 10^-2^ mM as per the timings in Figure 1B, immediately after TBI in larval fish and months prior to these behaviour recordings. **A-C)** Activity of an exemplar fish from the sham, TBI or TBI + LEV groups in early adulthood, showing TBI-treated fish spent less time spent near the novel object in the inner zone, and LEV partially recued this effect, represented as heatmaps of the circular arena. **D)** Inner zone preference in early adulthood (*P*< 0.0001) (Sham group n= 50, TBI n= 37, TBI + LEV n= 46). **E)** Inner zone preference in late adulthood (*P*= 0.0085) (Sham n= 32, Sham + LEV n= 21, TBI n= 22, TBI + LEV n= 18). Each data point represents one fish. The bar indicates the median of each group with 95% Cl. Significant differences between groups are indicated (*): *P*< .0332, (**): *P* < .0021, (***): *P* < .0002.

Behavioural deficits in the NOA test, induced by TBI, persisted at late adulthood. The TBI group continued to spend less time exploring the novel object in the inner zone compared to the sham groups (Figure 4E). However, the fish in the TBI + LEV group did not show a significant difference in object approach compared to the fish in the TBI group, highlighting a time-sensitive efficacy of LEV in reducing behavioural defects.

### 3.5 Sex-separated responses in OFT and NOA

Where available, data was analyzed by sex for both the open field (OFT) and novel object approach (NOA) tests. This analysis was conducted post-hoc to evaluate potential sex-dependent effects. No significant sex-dependent differences were observed in either behaviour (Figure 5A–F), but this could be due to the relatively small sample sizes. Thigmotaxis during the open field test in late adulthood showed no significance in the interaction of sex and treatment (F( 3,92)= 0.7031, P= 0.05526, Figure 5A) or between sex groups (*F* (3,92)= 0.5086, P= 0.6773, Figure 5A). However, there was a significant main effect of treatment *F*(1,92) = 0.7, *P* = .0178, Figure 5A), but post-hoc testing did not find any significance between groups. Similarly, inner zone preference during the NOA test was not significantly different between sexes (*F(*1,89) = 0.02495, P = 0.8748, Figure 5B). Distance moved in late adulthood did not differ by sex in either the OFT (*F*(1,537) = 1.301, P = .2546, Figure 5C) or NOA (*F*(1,90) = 0.05421, *P* = .8164, Figure 5D). Time spent immobile revealed a significant sex effect in the OFT and NOA only for the sham group (*F*(1,91) = 6.757, *P* = .0109, Figure 5E), (*F*(3,90) = 4.412, *P* = .1226, Figure 5F)

**Figure 5.**
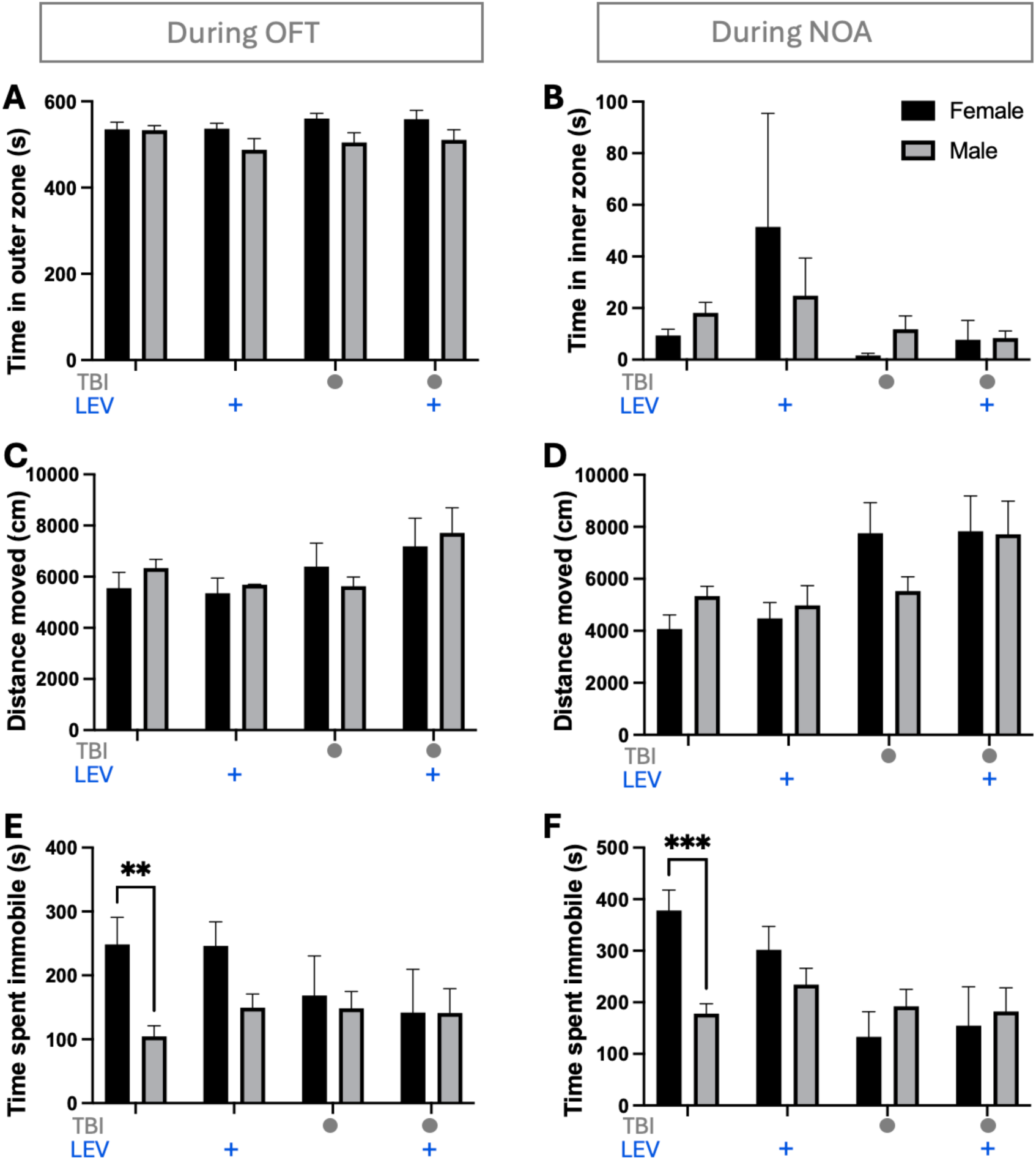
Sex-separated responses in open field and novel object approach test. Levetiracetam (LEV) was applied at 10^-2^ mM as per the timings in Figure 1B, immediately after TBI in larval fish and months prior to these behaviour recordings. **A)** Thigmotaxis during the open field test in late adulthood (*F*(7.000, 57.34) = 1.622, *P*= .1443). **B)** Inner zone preference during the NOA test in late adulthood (*P*= .0568). **C)** Total distance moved in OFT at late adulthood (*F*(7.000, 26.93) = 1.570, *P* = .1684). **D)** Total distance moved in NOA test at late adulthood (*P* = .0591). **E)** Time spent immobile during the OFT at late adulthood (*P* = *0*.0210). **F)** Time spent immobile during the NOA at late adulthood (*F*(7.000, 46.84) = 3.176, *P* = .0078). The bars indicate the mean of each group with SEM. Sample number in early adulthood: Sham n= 50, TBI n= 37, TBI + LEV n= 46). Sample number in late adulthood: Sham n= 32, Sham + LEV n= 21, TBI n= 22, TBI + LEV n= 18.

## 4. Discussion

This study aimed to determine whether a larval zebrafish model of TBI induces persistent neurobehavioral alterations detectable in adulthood and to evaluate whether post-injury treatment with LEV can mitigate these behavioural impairments. We found that our TBI model, which was administered at 3 dpf, did not negatively impact the survival of the fish (Figure 1C). There were no alterations in locomotion at each time point measured in both the OF and NOA, with the exception of LEV increasing locomotion in NOA test at late adulthood in TBI fish (Figure 2H). Taken together, the behavioural differences observed across assays are unlikely to be driven by physical injury or altered mobility. Our findings show that larval TBI leads to increased anxiety-like behaviour at early adulthood and induces a persistent avoidance of a novel object, the latter was rescued by LEV treatment (Figure 3 and 4). One sex-dependent variable was observed in the sham-treated fish where females spent more time immobile in both tests (Figure 5).

### 4.1 Larval TBI increased adult anxiety, an effect not measurably altered by LEV

Anxiety-like behaviour following larval TBI was first assessed using thigmotaxis in the OFT test, a well-established paradigm that measures increased wall-hugging behaviour as an indication of elevated baseline anxiety. Using this assay, anxiety-like behaviour was significantly elevated in TBI-exposed fish at early adulthood, and LEV treatment was unable to rescue this phenotype (Figure 3C). TBI cohorts, regardless of drug treatment, spent significantly more time near the arena walls compared to sham controls, indicating a persistent elevation in baseline anxiety at early adulthood. Notably, this effect was transient: by late adulthood, the sham cohort showed an elevation in the baseline anxiety levels, thus eliminating any observable anxiety differences at this stage. Older zebrafish (>12 months) have exhibited elevated anxiety in behavioural tests such as OFT and novel tank diving in other studies (Hudock & Kenney, 2024; Vasconcelos, Gordillo-Martinez, Ramos, & Lau, 2023).

A similar pattern emerged during the NOA test; however, the behavioural demands of this paradigm differ from those of the OFT. While the OFT primarily assesses generalized anxiety and exploratory behaviour in a novel environment, the NOA test considers fear of a novel stimulus, thus probing risk assessment, novelty processing, and the integration of sensory information with motivational and cognitive decision-making (Adolphs, 2013). The introduction of a novel object in the center of the arena therefore engages additional neural systems beyond those recruited during open field exploration alone. During early adulthood, TBI-exposed fish spent less time exploring the novel object and more time around the arena walls. No group differences were observed at late adulthood, again highlighting the baseline anxiety level increase corresponding to aging (Hudock & Kenney, 2024; Vasconcelos et al., 2023). Importantly, although behavioural variability was present across groups, the largest difference emerged between the sham + no drug and TBI + no drug groups when an object was introduced. This suggests that novelty amplified behavioural differences specifically in injured fish.

The heightened sensitivity to environmental novelty observed at early adulthood may therefore reflect altered sensory or contextual processing following TBI, rather than solely a generalized anxiety phenotype. Sensory hypersensitivity has been linked to concussive TBI and PTSD in humans (Hoffman, Lam, Hovda, Giza, & Fanselow, 2019), and similar mechanisms may underlie the exaggerated response to environmental change observed here. Together, these findings indicate that larval TBI produces fear-motivated behaviour that is not reversible by the LEV treatment we applied (Figure 3).

This newly documented anxiety at early adulthood, 8-10 months post-injury, demonstrates that the larval TBI paradigm developed by Alyenbaawi and colleagues can induce measurable long-lasting behavioural consequences. These results are consistent with previous studies that reported heightened anxiety following TBI in rodents (Hoffman et al., 2019) and humans (Hicks et al., 2021). Although antiepileptic drugs (AEDs) have been considered for the treatment of anxiety disorders previously (Van Ameringen, Mancini, Pipe, & Bennett, 2004), there has been limited research on TBI-induced anxiety models that have been treated with AEDs. While LEV did not seem to improve anxiety in TBI fish (Figure 3), further research into the effects of other AEDs on post-TBI behavioural deficits (akin to PTSD) and could prove to be significant.

### 4.2 LEV rescues persistent fear-related avoidance behaviour following TBI

The approach to a novel object indicates a tendency to explore with increased boldness relative to avoidance of the never-before seen stimulus. Untreated TBI fish spent significantly less time in proximity to the novel object compared to sham controls at both early and late adulthood, indicating sustained avoidance behaviour (Figure 4D & E). Given that locomotor activity was not measurably affected (Figure 2C & D), this avoidance cannot be attributed to impaired movement or reduced exploratory capacity due to physical limitations. Avoidance of requires complex interpretation because this behavioural state requires the integration of multiple cognitive processes, including threat evaluation, perception, and learning and memory. The persistent reluctance of TBI fish to approach a novel object could reflect alterations in fear processing or exploratory motivation. Impaired exploratory behaviour could be due to enhanced fear, as observed in concussed military personnel prone to PTSD (Glenn et al., 2017).

However, TBI can also alter sensory processing within subcortical sensory-emotional circuitry which may have also led to a reduction in exploratory behaviour as the fish cannot cognitively process the novel stimulus (Hoffman et al., 2019). Although the precise cognitive domain underlying this behavioural alteration cannot be definitively isolated, the persistent avoidance of a novel stimulus indicates a TBI-induced behavioural deficit that was partially rescued by LEV.

LEV treatment partially rescued TBI-induced behavioural deficits at early adulthood, as the Sham + no drug group and the TBI + LEV group spent significantly more time around the novel object than the TBI + no drug group (Figure 4D). At late adulthood, sham groups continued to show reduced approach to the novel object relative to both TBI and TBI + LEV groups, highlighting a time-sensitive effect of LEV in reducing behavioural deficits. These behaviour outcomes may reflect alterations within neural circuits governing fear-related responses. In humans, apprehension toward novelty involves structures like the amygdala, hippocampus, and prefrontal cortex (Tao, He, Lin, Liu, & Tao, 2021). Zebrafish also possess homologous brain regions that serve similar roles in processing emotions, memory, and executive functions (O’Connell & Hofmann, 2011). TBI may disrupt these conserved circuits, leading to dysregulated fear processing. LEV’s effects on synaptic transmission and neuronal excitability could have modulated activity within these fear-related networks, resulting in reduced apprehension and anxiety-like behaviours among injured fish.

### 4.3 Dissociating anxiety from cognitive fear processing

Importantly, the dissociation between normalized anxiety-like behaviour at late adulthood and persistent avoidance of the novel object suggests that the observed deficits in the Novel Object Approach test cannot be attributed solely to heightened anxiety. While TBI-induced anxiety-like behaviour was evident at early adulthood (increased time in the thigmotaxis zone in the OF), its resolution by late adulthood (no increased thigmotaxis preference in the OF) indicates that continued object avoidance is attributed to cognitive processing, such as fear learning, or exploratory motivation, rather than a generalized anxiogenic state.

Together, these results suggest that while AEDs such as LEV may effectively modulate certain post-TBI behavioural outcomes, their therapeutic effects need to be examined more thoroughly, with more doses and/or longer treatment exposures. Understanding how early-life brain injury differentially affects anxiety and fear circuitry is critical for developing targeted interventions for long-term neurobehavioural consequences of TBI.

## 5. Conclusion

Given the limited efficacy and safety of current clinical approaches for treating anxiety after TBI, there is a pressing need for experimental models that can both capture anxiety-related phenotypes and support systematic therapeutic discovery. The findings of the present study demonstrate that early-life TBI in zebrafish produces measurable anxiety-like behaviour in adulthood, providing suggesting that this injury model induces neurobehavioural alterations beyond molecular pathology. Importantly, the absence of gross locomotor impairments indicates that the observed behavioural changes are not attributable to physical limitations but rather reflect genuine behaviour dysfunctions. The validated anxiety phenotype established here positions this zebrafish model as a powerful platform for further drug screening. The experimental tractability and high-throughput capacity of larval zebrafish enable efficient evaluation of alternative pharmacological candidates, offering a scalable approach to identify novel treatments for TBI-induced anxiety that are currently lacking in clinical practice. While LEV was not effective as an anxiolytic, it was effective in reducing a fear-related response in zebrafish introduced to a novel object. Further cognitive tests should be conducted in addition to the NOA test to identify which cognitive domain is most likely impacted by TBI, inducing the behavioural deficits observed in this study.

Future works selecting an appropriate dose of larval TBI injury strength should trade-off survival, the robust capacity of the zebrafish brain to regenerate in early stages and through to adulthood, and the need to induce effect sizes that are robust and consistent enough to allow the impacts of treatment and sex to be detected. Selecting appropriate doses of pharmacology to apply immediately after TBI faces similar trade-offs. Whereas in larvae it is practical to apply various doses of intervention, e.g. five-to-ten doses can be used to probe the compounds’ in vivo hormesis and specificity and/or potential synergies, such approaches are less practical with longer experimental time-courses (Locskai et al., 2026). It would be of interest to determine if other doses (or time coursers) of LEV treatment are more efficacious in this paradigm, and whether that action could be improved by polypharmacy; indeed, our recent work demonstrates co-applying compounds (positive allosteric modulators that act on mGluR2 channels) can potentiate LEV’s therapeutic benefits by orders of magnitude, at least in the short term (Locskai et al., 2026).

## Acknowledgements

We are grateful to Melissa Kinley, Laszlo Locskai, and Andréa Johnson for discussions. Parastoo Razmara and Madison Oslund kindly provided critical feedback on an earlier draft of the writing. We appreciate the care of University of Alberta Animal Care staff and Veterinarians. SG was supported by the Natural Sciences and Engineering Research Council of Canada (NSERC) Canada Graduate Research Scholarship – Master’s (CGS-M) and the Walter H. Johns Graduate Fellowship. TCZ was supported by the Alberta Innovates Postdoctoral Recruitment Fellowship. Operating funds to WTA were in the form of donations from families who prefer to remain anonymous and to TJH as an NSERC Discovery Grant (03403).

